# Task-Level Value Affects Trial-Level Reward Processing

**DOI:** 10.1101/2021.09.16.460600

**Authors:** Cameron D. Hassall, Laurence T. Hunt, Clay B. Holroyd

## Abstract

Despite disagreement about how anterior cingulate cortex (ACC) supports decision making, a recent hypothesis suggests that activity in this region is best understood in the context of a task or series of tasks. One important task-level variable is *average reward* because it is both a known driver of effortful behaviour and an important determiner of the tasks in which we choose to engage. Here we asked how average task value affects reward-related ACC activity. To answer this question, we measured a reward-related signal said to be generated in ACC called the reward positivity (RewP) while participants gambled in three tasks of differing average value. The RewP was reduced in the high-value task, an effect that was not explainable by either reward magnitude or outcome expectancy. This result suggests that ACC does not evaluate outcomes and cues in isolation, but in the context of the value of the current task.

---

The role of anterior cingulate cortex (ACC) in decision making has been hotly debated (Holroyd & Verguts, 2021). According to one view, ACC supports reward-guided behaviours like foraging by encoding the value of switching from one task to another (Kolling et al., 2016). Others argue that ACC selects from among competing tasks by computing the *expected value of control*, balancing effort and task value (EVC: Shenhav et al., 2013). Computationally, ACC has been proposed to follow the principles of hierarchical reinforcement learning, exerting control over lower-level systems when ongoing rewards are worse than the average task value (Holroyd & McClure, 2015).

In each of these approaches, ACC activity depends on task value. This claim has empirical support; neurons in monkey ACC, for example, encode both trial-by-trial reward and average task value (Amiez et al., 2006). But how does *trial*-level reward processing interact with *task*-level value processing in ACC? One possibility is via a reward prediction error (RPE), a concept borrowed from reinforcement learning (RL) that indicates whether ongoing events are better or worse than expected (Sutton & Barto, 2018). RPEs are proposed to be carried by the midbrain dopamine system to striatal and cortical targets (Schultz et al., 1997), including ACC (Amiez et al., 2005; Holroyd & Coles, 2002). According to the standard “model-free” RL approach, called temporal difference learning, the RPE elicited by reward delivery is computed by subtracting the expected value of an action from the just-received reward. The standard reinforcement learning view is that RPEs are “one-step” computations, comparing the value of the current state to the value of the previous state only. Under this view, dopaminergic RPEs depend only on action/reward values, not the average task value.

Alternatively, midbrain dopamine activity may index a more general RPE rather than just an immediate, “one-step” RPE. Indeed, there is evidence that midbrain RPEs are sensitive to task context. For example, monkey dopamine neurons appear to track patterns in reward history beyond what would be predicted by a standard RL algorithm (Nakahara et al., 2004). Reinforcement learning algorithms have therefore been augmented with internal models of the environment resulting in “model-based” RL algorithms. Both model-free and model-based reinforcement learning computations appear to drive activity of midbrain dopamine neurons (Collins & Cockburn, 2020; Daw et al., 2011). There is therefore good reason to believe that midbrain RPE signals may be sensitive to task-level factors such as average task value.

In humans, midbrain RPE signals are thought to modulate ACC activity in a way that is measurable at the scalp. It has been proposed that a component of the event-related potential (ERP) called the reward positivity (RewP) varies as a function of RPE magnitude (Holroyd & Coles, 2002; Sambrook & Goslin, 2015). In other words, the RewP provides a convenient readout of the degree to which outcomes differ from learned *action* values. Although there is debate about the neural source of the RewP, many studies have localized it to ACC (see review by Walsh & Anderson, 2012). Our goal here was to investigate whether and how reward-related ACC activity, as indexed by the RewP, varies with *task* value. Importantly, action value and task value are partially dissociable in reinforcement learning frameworks: High-value actions can occur in low-value tasks, and vice-versa.

To answer this question, we manipulated task value and action value by varying the proportion of “low-value” and “high-value” actions in three probabilistic learning tasks. EEG was recorded from participants as they attempted to learn correct actions for six predictive cues. Some of the cues were “high-value”, indicating that a correct response would likely yield a reward. Other cues were “low-value”, indicating that reward and non-reward outcomes were equally likely regardless of the response. We did not vary the value of the reward itself, which can affect the amplitude of the RewP (Kreussel et al., 2012; Sambrook & Goslin, 2015). Rather, we varied the proportion of high- and low-value cues in each task: either all low-value, all high-value, or an even split. Our goal here was to vary reward at the task level while keeping trial-level rewards constant. Importantly, the same cue type (same reward expectancy and reward magnitude) appeared in multiple tasks, allowing us to isolate the effect of task value on feedback-locked signals.

In addition to the RewP, we also examined the cue-locked ERP. There is evidence that reward-predicting cues can elicit a RewP-like signal (Holroyd et al., 2011; Krigolson et al., 2014). The computational explanation for this is that RPEs propagate backward in time to the earliest indicator that things are better or worse than expected. In a task with mixed high- and low-value cues, we might therefore expect high-value cues to elicit a positive prediction error (a positive RewP deflection) relative to low-value cues. Conversely, we would not expect a cue-locked prediction error to be elicited in a task with uniform cue values, because the cues all make the same prediction about upcoming rewards within the task. To summarize, trial cues ought to elicit positive/negative prediction errors in the “mixed value” task and no prediction error in the “uniform value” tasks

## Method

### Participants

We tested 36 participants with no known neurological impairments and with normal or corrected-to-normal vision (11 male, 5 left-handed, 1 ambidextrous) across two testing sites. At the first testing site we tested 24 University of Victoria (UVic) undergraduate participants. At the second testing site we tested 12 University of Oxford (Oxford) graduate students and community members. Participant ages ranged from 18 to 77, *M* = 27.6, *SD* = 14.5. UVic participants received bonus credit in an undergraduate psychology course. Oxford participants were compensated £20 (£10 per hour). Additionally, each participant received a performance-dependent monetary bonus. The study was approved by the University of Victoria Human Research Ethics Board and the Medical Sciences Interdivisional Research Ethics Committee at the University of Oxford. All participants gave written informed consent.

This experiment required that participants learn to make optimal responses when this was possible, i.e., in response to high-value cues. As such, we applied the following a priori criterion: only the data of participants who made a correct response on at least 60% of the learnable trials in both the mid-value task and the high-value task were included in the main EEG analysis. 12 of the 36 participants did not meet this criterion and their data were therefore removed from the main analysis. However, the data of these 12 participants were included in a correlational analysis relating performance to the RewP. Finally, the data of one participant were excluded from all analyses due to excessive EEG artifacts (across all conditions, the average trial rejection rate for this participant was 87%). This left a total of 24 participants for the main analysis and 35 participants for the correlational analysis.

### Apparatus and Procedure

Participants were seated 60 cm in front of an LCD display (60 Hz, 1024 by 1280 pixels). Visual stimuli were presented using the Psychophysics Toolbox Extension (Brainard, 1997; Kleiner et al., 2007; Pelli, 1997) for MATLAB (Version 8.2, MathWorks, Natick, USA). Participants were given both verbal and written instructions in which they were asked to minimize head and eye movements.

Participants were told that they would be gambling in three different casinos. Each casino contained six different slot machines, and each slot machine was represented by a unique, coloured shape. Prior to “entering” a casino, the message “New Casino, New Coloured Shapes” was displayed. Shapes were reused across casinos, but with different colours (randomly chosen for each participant). The slot machines were described to participants as having two arms – a left arm and a right arm. Participants were instructed to, upon the appearance of a slot machine (a coloured shape), select one of the arms by pressing the corresponding key on a keyboard (the s-key to select the left arm, or the k-key to select the right arm). Participants were also told that their gamble would result in win or a loss – each outcome represented by a randomly-assigned fruit – and that for each slot machine “pulling one arm may be more likely to result in a win compared to pulling the other arm”. Wins resulted in a gain of $0.03, while losses resulted in no gain ($0.00). Participants were informed that their goal was to win as much money as possible and that they would be paid their total at the end of the experiment.

Trials were grouped by casino, and each casino was entered only once. Casino order was counterbalanced with six possible casino orderings and four participants assigned to each ordering. Within a casino, participants encountered the 6 slot machines 24 times in random order (144 gambles). Unknown to participants, there were two types of slot machine: high-value and low-value. Each high-value slot machine had one arm (randomly chosen) that, when selected, would result in a win with 80% probability (selecting the other arm resulted in a win with a 20% probability). In contrast, low-value slot machines had no correct answer: there was a 50% probability of winning, regardless of which arm was pulled. The casinos contained either only low-value slot machines (the low-value task), high-value slot machines (the high-value task), or an even split between low- and high-value slot machines (the mid-value task). As each slot machine was encountered the same number of times, there was no learning advantage related to the number of exposures. Upon leaving a casino the total amount won within that casino was displayed.

Each trial began with the appearance of a white fixation cross presented against a black background (Figure 1). The fixation cross, and all other visual stimuli, subtended 2° of visual angle. After 400–600 ms, the fixation cross was replaced by the coloured shape representing the current trial’s slot machine (the “cue”). After 1000 ms, a 50 ms 400 Hz sine tone signalled participants to choose an arm (left or right) by pressing the corresponding key on a keyboard. The purpose of the delay between the cue and the tone was to isolate cue-related neural activity from response-related neural activity. The coloured shape/slot machine remained on the display until the participant responded (or until 2000 ms if no valid response was made). Finally, another fixation cross appeared for 400–600 ms, followed by the feedback stimulus for 1000 ms. Two images of fruit were used as feedback stimuli, chosen at random for each participant from six possible images. If the participant responded prior to the onset of the tone, the message “too fast” was displayed instead. If no response was made within the 2000 ms response window, or if a non-response key was pressed, the message “invalid” was displayed. In both cases (fast/invalid responses) the trial was excluded from both the behavioural analysis and the EEG analysis.

**Figure 1.**
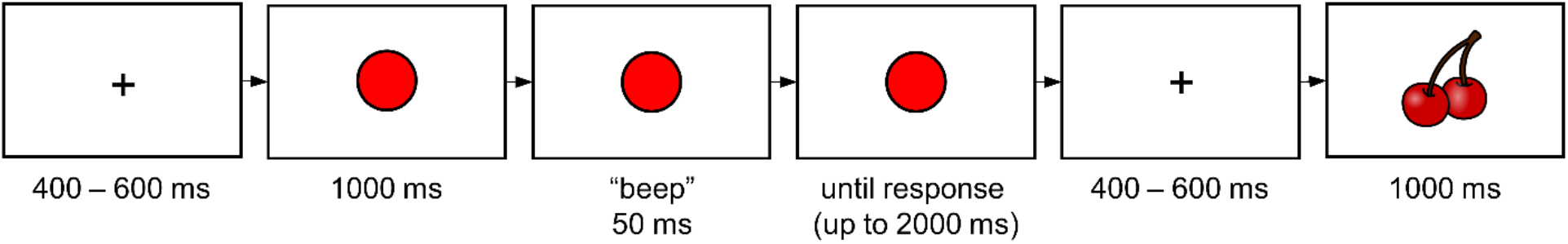
Task overview. After making a left or right button press, participants were shown fruit indicating the outcome (win or loss).

### Data Collection

For each trial, the experimental software recorded reaction time (time since the onset of the tone), response (left, right, invalid, or no response), and casino/slot machine information. We also recorded casino value (low, mid, or high), slot machine value (low or high), trial outcome (win or lose) and – for high-value slot machines – whether or not the correct arm was chosen (that is, the arm more likely to result in a win).

EEG was recorded from 35 Ag/AgCl electrode locations using Brain Vision Recorder (Brain Products GmbH, Gilching, Germany). The electrodes were mounted in a fitted cap (EASYCAP GmbH, Wörthsee, Germany) with a standard 10-20 layout and were recorded with respect to an average reference. Electrode impedances were lowered below 20 kΩ prior to recording and the EEG data were sampled at 250 Hz and amplified (UVic: Quick Amp, Oxford: actiCHamp Plus, Brain Products GmbH, Gilching, Germany).

## Data Analysis

For all three tasks (low-value, mid-value, high-value) we computed the mean proportion of trials resulting in a win. For the mid-value and high-value tasks we computed, for each participant and task, the likelihood of a correct response to the high-value cues. (Recall that low-value cues had no correct response.) As mentioned previously, participants for whom this likelihood fell below 60%, in either the mid-value task or the high-value task, were excluded from the main analyses (seven participants in total). For visualization purposes, we computed a sliding window measure of performance in the mid-value and high-value tasks: the proportion of wins in 30 trials, moving the window by 1 trial each time.

The EEG was analyzed in MATLAB 2022a (MathWorks, Natick, USA) using the EEGLAB library (Delorme & Makeig, 2004). EEG data were first bandpass filtered (0.1– 30 Hz, UVic: 60 Hz notch, Oxford: 50 Hz notch) and re-referenced to the average of the mastoid signals. Ocular artifacts were identified and removed using independent component analysis (ICA). The ICA was trained on three-second epoch starting at the presentation of the cue. Ocular components were identified using the *iclabel* function and removed from the continuous data. We removed components which *iclabel* determined were more likely to be eye-related than brain-related.

### ERP Analysis

We constructed 800 ms epochs around the appearance of each cue and feedback stimulus (−200 to 600 ms). The data were then checked for remaining artifacts, and any epochs containing a change in voltage of more than 40 µV per sample point or an overall change in voltage of more than 150 µV across the epoch were excluded from the analysis. On average, 1.0% (*SD* = 1.5%) of cue-locked epochs and 0.8% (*SD* = 1.3%) of feedback-locked epochs were excluded. We also excluded the first ten trials of each task, as participants became familiarized with the new cues. We then averaged over the remaining epochs to create mean cue-locked, win-locked, and loss-locked waveforms for each task (low-value, mid-value, high-value), cue (low-value, high-value), and participant. To quantify the RewP, we used the difference wave method by subtracting the mean loss ERP from the corresponding mean win ERP for each task and cue. The difference wave approach is especially useful for isolating the RewP because it minimizes component overlap (Krigolson, 2017; Luck, 2014). We then examined the electrode location and time window recommended by Sambrook and Goslin (2015): FCz from 240–340 ms post feedback. The RewP was defined as the maximum voltage recorded within this window for each task value, cue value, and participant. For completeness, we also computed the mean voltage from 240–340 ms at electrode FCz for each cue value (low, high), outcome (win, loss), and task value (low, mid, high). We then computed effect sizes (Cohen’s *d*) comparing similar cue values and outcomes (see Supplemental Table 1). A similar approach was used for the cue-locked ERPs (see Supplemental Material).

### Inferential Statistics

To confirm our task value manipulation, we compared the mean proportion of wins in each task using a one-way repeated-measures ANOVA. Partial and generalized eta-squared were computed as:

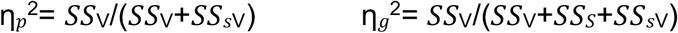

where *SS*_V_ is the sum of squares of the value effect (low, mid, high), *SS*_sV_ is the error sum of squares of the value effect, and *SS*_S_ is the sum of squares between subjects. To test whether performance differed between the mid-value and high-value block – that is, whether participants learned about high-value cues differently – we used a repeated-measures *t*-test.

To determine the effect of average task value on the RewP, we compared RewP scores using repeated measures *t*-tests. To avoid possible outcome frequency confounds, we only compared RewP scores from conditions with similar outcome frequencies (Krigolson, 2017). Specifically, we compared the low-task, low-cue RewP (infrequent rewards in an infrequent reward context) to the mid-task, low-cue RewP (infrequent rewards in a mid-frequency reward context). We then compared the mid-task, high-cue RewP (frequent rewards in a mid-frequency reward context) to the high-task, high-cue RewP (frequent rewards in a frequent reward context).

For all *t*-tests, we first checked the assumption of normality of each variable using the Shapiro-Wilk test. The assumption was met for all variables except for the RewP score in the high-task, high-cue condition. As the *t*-test is robust to non-normality, no correction was made. For each comparison, we also computed Cohen’s *d* for paired-samples t-tests as:

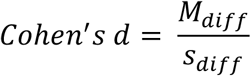

where *M*_*diff*_ is the mean difference between the scores being compared and *s*_*diff*_ is the standard deviation of the difference of the scores being compared (Cumming, 2014).

### Modelling

We used computational modelling to explore the possibility that participants may have used a different strategy in each task, either due to differences in average task value, or in response to the different outcome probabilities. For example, participants may have adjusted the degree to which they weighed past outcomes when considering their current choice. Participants may have also adjusted their rate of exploration. Each of these behaviours (weighing of past outcomes, exploration) is expressed in a reinforcement learning (RL) model as the learning rate and temperature parameters, respectively. To determine whether these behaviours differed in our tasks, we modelled trial-by-trial choices for each participant, task value, and cue value.

For each cue, the model maintained action values *Q* associated with each response (left, right). Following a chosen action *a*, a prediction error *δ* was computed according to:

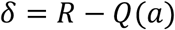

The action value of the chosen action *a* was then updated according to:

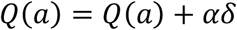

where *α* is the learning rate. The likelihood of a particular action *a* was determined by the *Q*-values according to the softmax equation, depending on the temperature parameter *τ*:

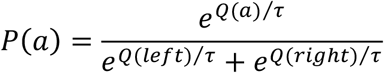

Model fit was assessed by combining the chosen action probabilities across all trials *t* using negative log-likelihood:

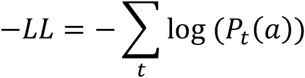

A good model fit meant that the model assigned a high likelihood to the chosen action on each trial, resulting in a low negative log-likelihood. Parameters (*α, τ*) were individually fit for each participant, task value, and cue value to minimize negative log-likelihood using the MATLAB function *fmincon* (Optimization Toolbox, Version 9.3, MathWorks, Natick, USA). Parameters were restricted to recoverable values: learning rates from 0.1 to 1 and temperatures from 0.01 to 1 (see Supplemental Material for parameter and model recovery). For comparison, we also computed the negative log-likelihood of a random model, which used an action likelihood of 0.5 on each trial. Model fits for each cue value were then compared via repeated-measures *t*-tests, as described above. We also tested whether the optimized model parameters (*α, τ*) differed by cue value.

## Results

### Behavioural Results

The proportion of winning trials differed by task (low-value: 49.82 %, 95% CI [48.33, 51.31], mid-value: 59.07 %, 95% CI [57.24, 60.90], high-value: 67.53 %, 95% CI [64.61, 70.46]), *F*(2,46) = 69.38, *p* < .001, η_p_^2^ = 0.75, η_g_^2^ = 0.67. Note that since each win was worth the same amount, the total reward also differed between tasks (Figure 2a). There was no evidence of a difference between mean performance in the mid-value block (81.11%, 95% CI [76.26, 85.96]) and mean performance in the high-value block (78.06%, 95% CI [73.64, 82.48]), *t*(23) = 1.29, *p* = .211, Cohen’s *d* = 0.21 (Figure 2b–c). Recall that mean performance for both the mid-value blocks and the high-value blocks was defined as the proportion of high-value cues that were responded to correctly.

**Figure 2.**
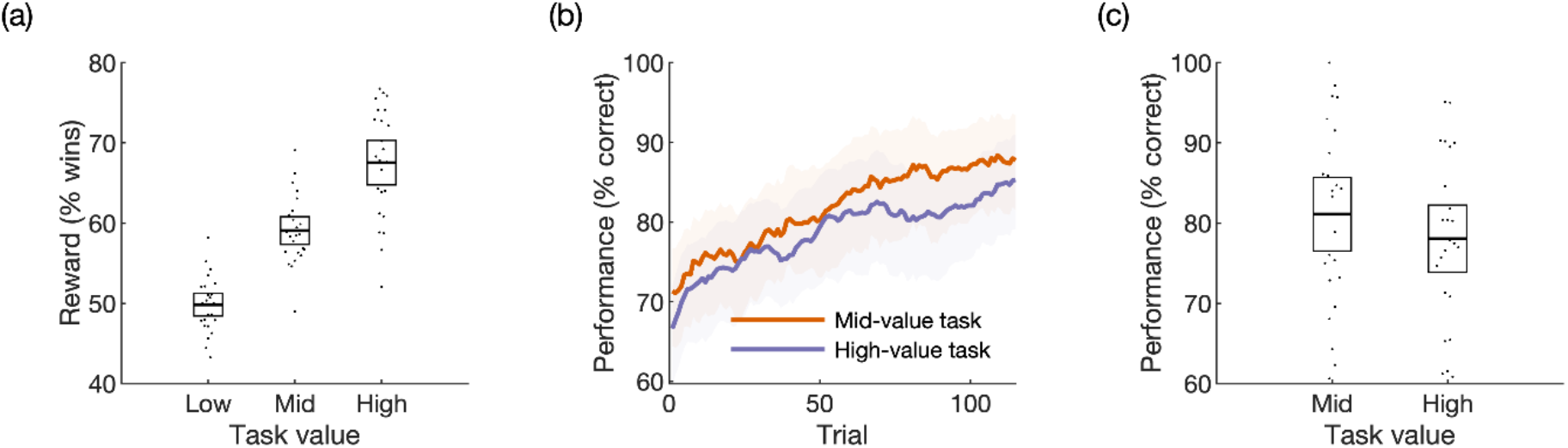
Behavioural results. (a) Mean proportion of wins in each task. (b) Performance curves in the tasks where correct responses were possible (mid-value and high-value tasks), indicating improvement over time. (c) Mean proportion of correct responses within the mid-value and high-value tasks. The shaded regions in each plot indicate 95% confidence intervals. Dots correspond to individual participants.

### Feedback-Locked EEG (RewP)

After collapsing across the average feedback-locked waveforms (Figure 3a) we observed a scalp topography difference consistent with a RewP (Figure 3b). After constructing conditional difference waves (Figure 3c), we observed a significant effect of task value on the RewP amplitude elicited by high-value cues, indicating that RewP size was modulated by task value: The RewP following high-value cues was larger in the mid-value task (7.39 µV, 95% CI [5.30, 9.48]), *t*(23) = 2.53, *p* = .019, Cohen’s *d* = 0.52 than in the high-value task (5.29 µV, 95% CI [3.83, 6.75]). The same difference was not observed when the RewP following low-value cues in the low-value task (6.50, 95% CI [4.73, 8.27]) was compared to the RewP following low-value cues in the mid-value task (6.98, 95% CI [4.88, 9.08], *t*(23) = 0.51, *p* = .615, Cohen’s *d* = 0.10 (Figure 3d). The observed RewP increase (mid-value task minus high-value task) depended on the mean performance across all conditions, *r* = 0.45, *p* = .007. In other words, more accurate participants showed a greater RewP increase in the mid-value task (Figure 4).

**Figure 3.**
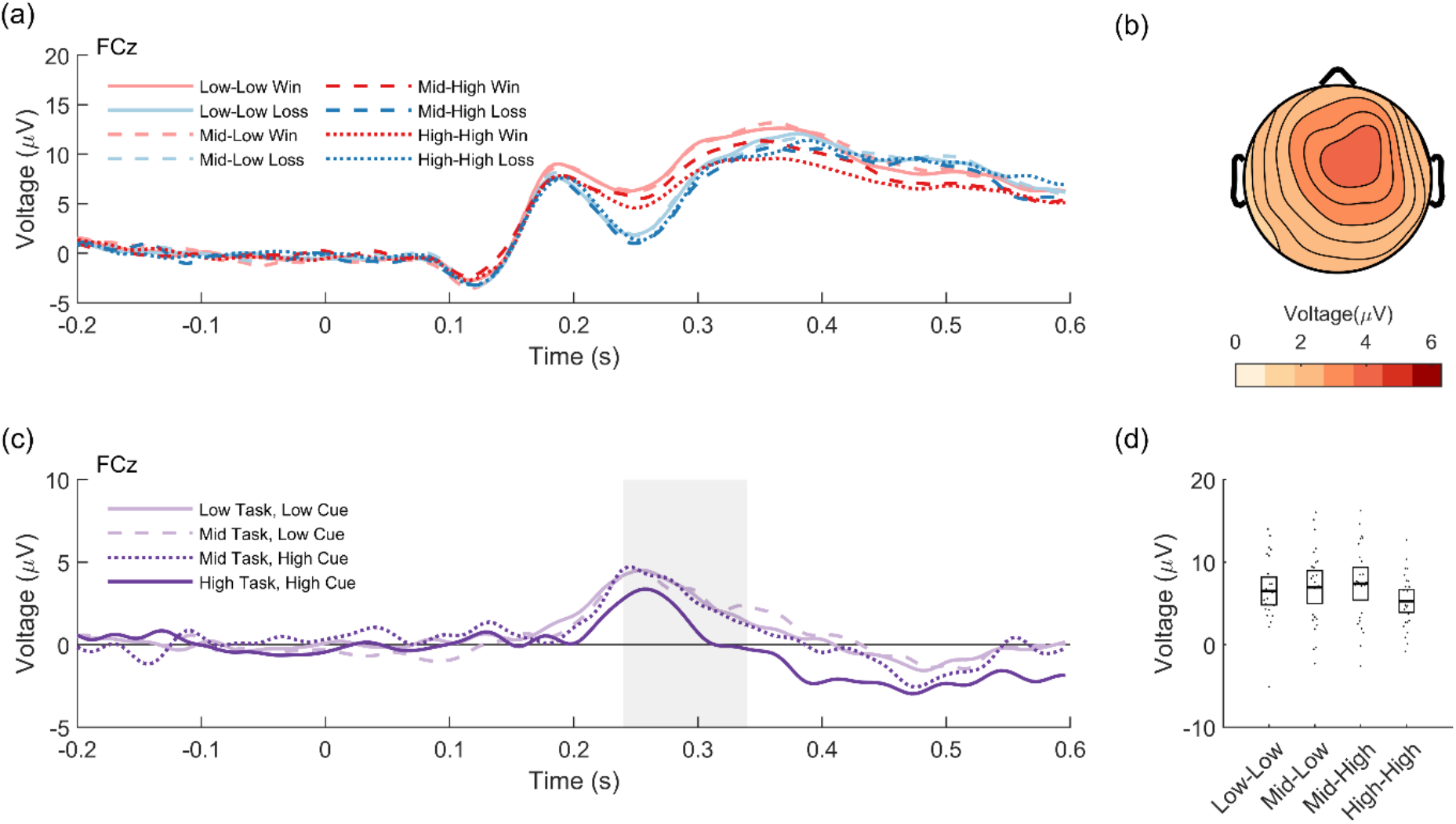
The reward positivity is reduced when task value is high. (a) Win/loss waveforms by task value (low, mid, high) and cue value (low, high). (b) Scalp distribution of the win-loss difference collapsed across conditions. (c) Difference waveforms by task value (low, mid, high) and cue value (low, high). The shaded area shows the region of analysis. (d) RewP scores by task value (low, mid, high) and cue value (low, high). Waveforms are time-locked to feedback onset. Error bars indicate 95% confidence intervals. Dots correspond to individual participants.

**Figure 4.**
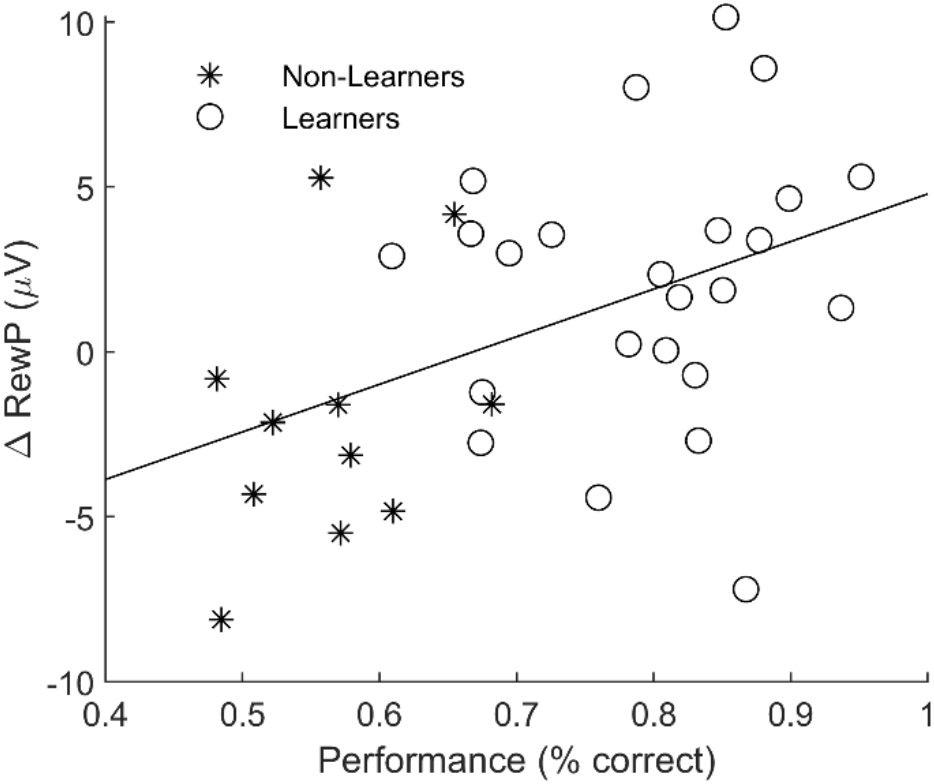
The reward positivity effect depends on performance. The difference in RewP between the mid-value and high-value tasks correlated across participants positively with task performance (greater difference associated with more accurate performance).

### Cue-Locked EEG (RewP)

No cue-locked RewP difference was observed (see Supplemental Material).

### Modelling Results

In the mid-value task, the fit learning rate parameter *α* was greater for low-value cues (0.52, 95% CI [0.39, 0.65]) than high-value cues (0.23, 95% CI [0.14, 0.32]), *t*(23) = 3.72, *p* = .001, Cohen’s *d* = 0.76, suggesting that participants weighed recent outcomes more heavily for low-value cues. The same difference was not observed between low-value cues in the low-value task (0.46, 95% CI [0.32, 0.60]) and high-value cues in the high-value task (0.32, 95% CI [0.20, 0.44]), *t*(23) = 1.64, *p* = .114, Cohen’s *d* = 0.34. See Figure 5a.

**Figure 5.**
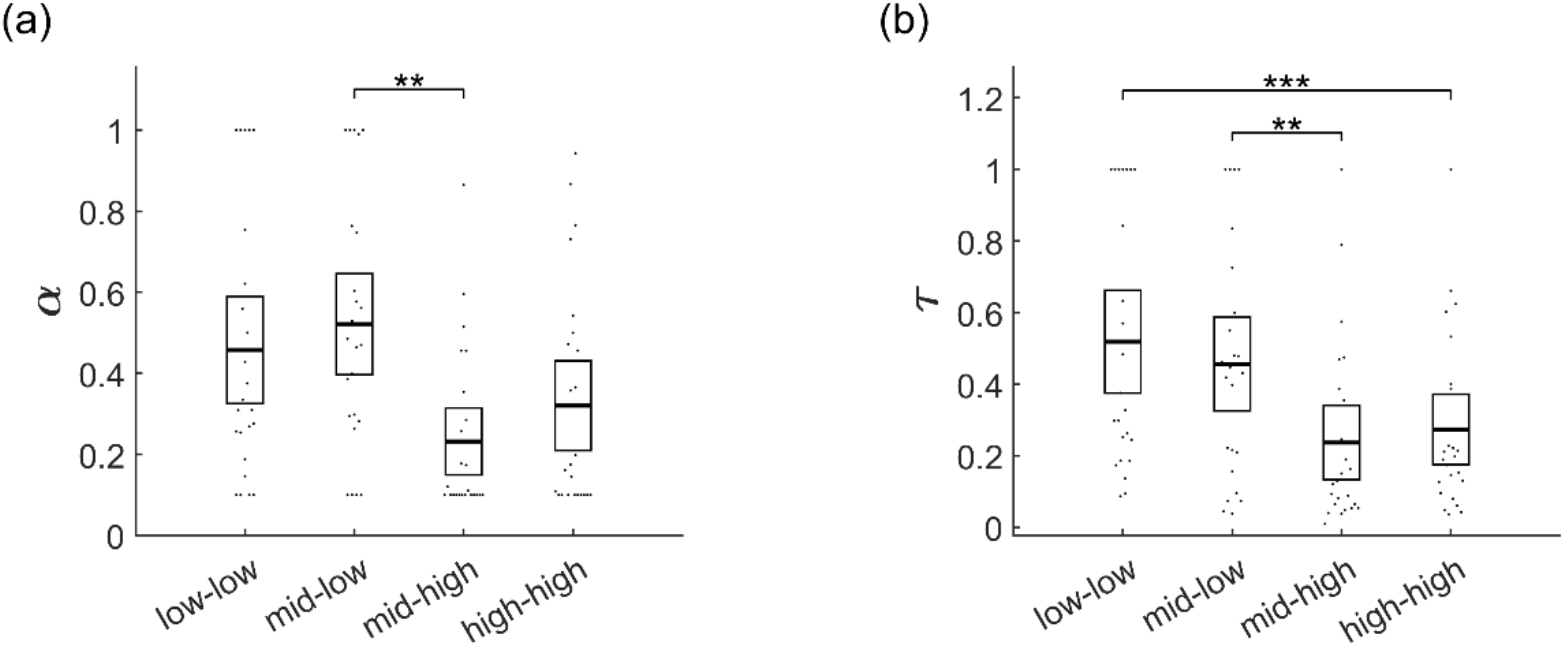
Computational modelling suggests participants used a different strategy for low-value cues. (a) The learning rate parameter, α, fit to each participant, was greater for mid-low cues compared to mid-high cues. (b) The temperature parameter, *τ*, fit to each participant, was greater for low-value cues compared to high-value cues. Dots correspond to individual participants.

Furthermore, within the mid-value task, *τ* was greater for low-value cues (0.46, 95% CI [0.32, 0.59]) compared to high-value cues (0.24, 95% CI [0.13, 0.35]), *t*(23) = 3.13, *p* = .005, Cohen’s *d* = 0.64 (Figure 5b). Likewise, the fit temperature parameter *τ* was greater in the low-value task (0.52, 95% CI [0.37, 0.67]) compared to the high-value task (0.27, 95% CI [0.17, 0.38]), *t*(24) = 3.79, *p* < .001, Cohen’s *d* = 0.77, suggesting that choices were more random and less value-driven when the average task value was low. See Supplemental Material for parameter and model recovery results.

## Discussion

ACC activity is often studied by focussing on relatively short-term, within-trial processing of actions and events, despite empirical evidence and computational considerations suggesting that this region is more concerned about global aspects of task performance (Holroyd & McClure, 2015; Holroyd & Verguts, 2021; Holroyd & Yeung, 2012; Umemoto et al., 2017). Here we show evidence that trial-to-trial ACC responses, as indexed by the RewP, are influenced by task value. We measured these signals in three tasks of varying average value. Despite matching outcomes for expectancy and trial-level value, the RewP was reduced when average task value was high.

This result is in line with a growing body of literature highlighting the importance of task context in understanding ACC activity. By “context” we mean task value (as in the present study) and other relevant task-level variables. For example, RewP amplitude is sensitive to the range of possible outcomes indicated by task instructions, not to the objective values of these outcomes (Holroyd et al., 2004). Task-level goals also matter: the amplitude of the RewP is greater when reward indicates progress towards a goal (Osinsky et al., 2017; Shahnazian et al., 2018). The RewP is also enhanced for self-relevant rewards (Krigolson et al., 2013; Yu & Zhou, 2006) and in agentic situations, when perceived control of outcomes is high (Hassall et al., 2019; Martin & Potts, 2011; Mühlberger et al., 2017; Yeung et al., 2005).

Traditional RL models learn action values without representing task context (such as average task value) explicitly. However, here we controlled for trial-level expectancy and yet observed a task-level influence on the RewP. This result – a reduced RewP when task value is high – is inconsistent with a model that learns at the level of actions only, but may be consistent with an average-reward learning model in which prediction errors are defined as the difference between trial rewards and the average task value (e.g., Holroyd & McClure, 2015). Under this view, the RewP ought to be reduced when trial outcomes are closer in value to the average task value, as we observed for outcomes following high-value cues in the high-value task. Note that we did not observe a similar effect for low-value outcomes (outcomes following low-value cues). This result may be due to a difference in strategy; participants used a larger learning rate when attempting to learn about low-value cues, suggesting they were weighing recent outcomes more heavily compared to high-value cues. Additionally, low-value cues were associated with a larger temperature parameter, indicative of enhanced exploration for the more difficult problems.

There is empirical evidence that the RewP is sensitive to average reward. For example, we have previously observed that a simple RL model fails to account for RewP amplitude when participants play virtual casino games in virtual casinos (Umemoto et al., 2017). There, a RewP was observed to casino game outcomes and to casinos themselves – in other words, ACC is sensitive to both game values and casino values. To explain these neural responses computationally, it was necessary to combine a simple RL model with an average-value learning model. In general, representing contextual information like average task value can be beneficial in a hierarchical framework in which a limited resource such as cognitive control needs to be regulated across multiple levels. These features are implemented in the hierarchical reinforcement learning (HRL) model of ACC (Holroyd & McClure, 2015). Under this framework, ACC signals a need for control when outcomes are worse than the average task value.

Besides the RewP, there is other evidence that the brain tracks average reward over time. In the short-term, unexpected rewards (RPEs) elicit phasic midbrain dopaminergic activity that is linked to trial-to-trial improvements in behaviour (Pessiglione et al., 2006). Subjectively, RPEs are associated with momentary changes in happiness (Rutledge et al., 2014, 2015). However, the brain also tracks rewards in the long-term. There is fMRI evidence that BOLD activity in midbrain correlates with average reward, even when controlling for RPEs (Rigoli et al., 2016). The proposed mechanism behind this effect is tonic midbrain dopamine, which is thought to track average reward rate in order to support motivational vigour (Niv et al., 2007).

Relevant here is the possibility that phasic and tonic dopamine interact. In particular, a high level of tonic dopamine is thought to suppress phasic activity (Bilder et al., 2004; Grace, 1991, 2000). One piece of evidence for this comes from EEG; the RewP, which is linked to phasic dopamine, is reduced in individuals with increased tonic dopamine (Marco-Pallarés et al., 2009; Mueller et al., 2014; but see Heitland et al., 2012; Foti & Hajcak, 2012). This result suggests that the interaction between phasic and tonic dopamine may provide a dopamine-based mechanism for our ERP result: increased tonic activity in the high-value task relative to the mid-value task resulted in reduced phasic activity (and a concomitant decrease in RewP amplitude).

The ACC has long been known to respond to trial-level events such as cues and rewards (Holroyd & Coles, 2002). Another picture of the ACC is emerging – that of a region tasked with supporting extended sequences of actions (Holroyd & Verguts, 2021; Holroyd & Yeung, 2012). Here we provide evidence that ACC activity depends not only on trial-level features (is this a good outcome/cue?) but also task-level variables (how good is the task?) We suggest that a global view of ACC function will prove more fruitful than studying its neural responses at a molecular level.

## Data Availability Statement

Data for participants at the Oxford testing site is available at https://doi.org/10.18112/openneuro.ds004147.v1.0.0. Participants at the UVic testing site did not consent for their raw or preprocessed EEG data files to be publicly shared. Task and analysis scripts are available at https://github.com/chassall/averagetaskvalue.

## Acknowledgements

C.D.H. was supported by a Natural Sciences and Engineering Research Council of Canada (NSERC) Postdoctoral Fellowship (PDF-546078-2020). L.T.H. was supported by a Sir Henry Dale Fellowship from the Royal Society and Wellcome (208789/Z/17/Z). C.B.H. was supported by Natural Sciences and Engineering Research Council of Canada (NSERC) Grant # 312409-05 and by funding from the European Research Council (ERC) under the EU’s Horizon 2020 Research and Innovation Programme (grant agreement no. 787307). Data collection at the Oxford testing site was supported by the NIHR Oxford Health Biomedical Research Centre. The Oxford testing site is part of the Wellcome Centre for Integrative Neuroimaging, which was supported by core funding from Wellcome Trust (203139/Z/16/Z).

