## Supplemental Material for "Task-Level Value Affects Trial-Level Reward Processing"

**Supplementary Material**

### Cue-Locked ERP Analysis

As with our feedback-locked analysis, we examined the average epoched EEG at electrode FCz from 240–340 ms post cue. ERP scores were generated by computing the average voltage in this window for each condition. We then analyzed the resulting ERP scores using a one-way repeated-measures ANOVA. No cue-locked difference was observed at FCz, $F(3,69) = 0.34$, $p = .80$, $\eta_p^2 = 0.01$, $\eta_g^2 = 0.001$.

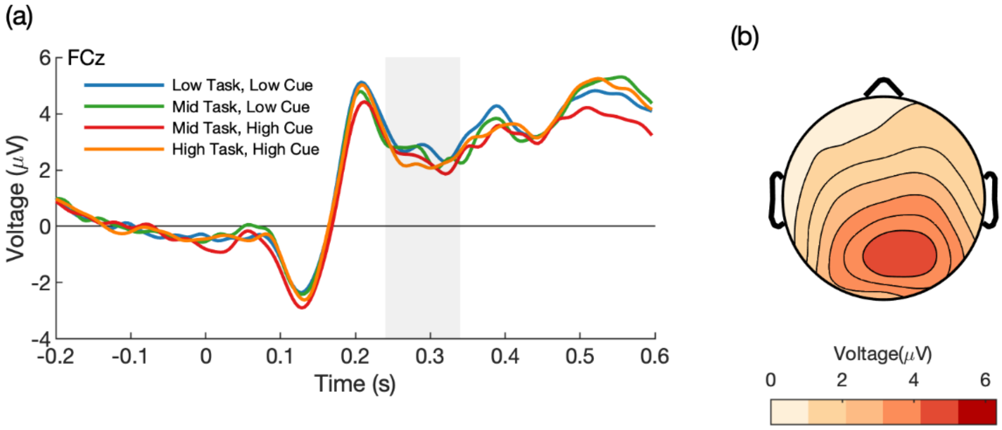


*Supplementary Figure 1.* **No cue-locked RewP.** (a) Average waveforms for each condition. The shaded region shows the analysis window. (b) Scalp topography of the average EEG from 240-340 ms post cue (all conditions combined).

Supplemental Table 1

*Mean voltages and effect sizes when comparing wins only or losses only*

| | | Task Value | | | |
|---|---|---|---|---|---|
| Cue Value | Outcome | Low (μV) | Mid (μV) | High (μV) | Effect Size (Cohen's *d*) |
| Low | Win | 9.50 | 9.44 | | -0.01 |
| | Loss | 6.32 | 6.18 | | -0.03 |
| High | Win | | 8.41 | 7.22 | 0.27 |
| | Loss | | 5.33 | 5.66 | -0.07 |

### Modelled Parameters, Parameter Recovery, and Model Recovery

In each task, a reinforcement learning model explained choice behaviour better than a random baseline model. However, we observed that model fit was better (lower -*LL*) in the high-value task (57.91, 95% CI [49.09, 66.73]) compared to the low-value task (81.89, 95% CI [77.78, 86.00]), $t(23) = 6.05$, $p < .001$, Cohen's $d = 1.23$. Similarly, within the mid-value task the fit for high-value cues (25.48, 95% CI [19.84, 31.12]) was better compared to the fit for low-value cues (37.24, 95% CI [33.60, 40.87]), $t(23) = 4.21$, $p < .001$, Cohen's $d = 0.86$. Note that there were half as many mid-low and mid-high trials compared to low-low and low-high trials. For visualization purposes we doubled the model fits for mid-low and mid-high cues (Supplementary Figure 2).

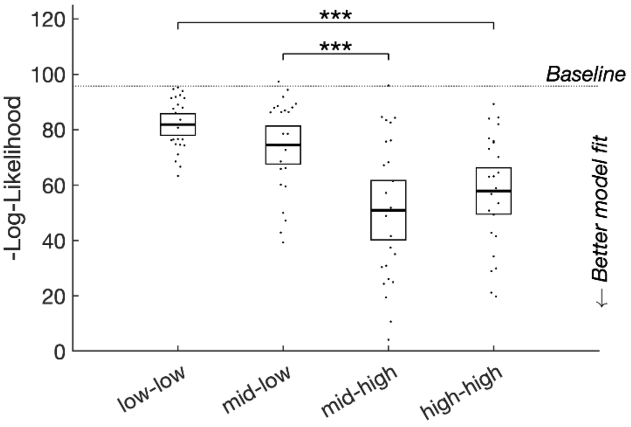


*Supplementary Figure 2.* **Modelled fits.** Compared to baseline (random choices), behaviour could be better explained by a reinforcement learning (RL) model. However, model fit was significantly worse in the low-value task compared to the high-value task and (in the mid-value task) for low-value cues compared to high-value cues.

#### *Parameter Recovery*

To assess parameter recovery, we simulated responses for 100 participants using a range of learning rates (0.01, 0.1, 0.2, 0.3, 0.4, 0.5, 0.6, 0.7, 0.8, 0.9, 1) and temperatures (0.01, 0.1, 0.2, 0.3, 0.4, 0.5, 0.6, 0.7, 0.8, 0.9, 1). For each parameter combination we ran the model-fitting procedure described in the main manuscript. We then examined the recovered parameters in each task. There was a good level of agreement between the original parameters and the recovered parameters (Supplementary Figure 3).

### ***Model Recovery***

For each simulated data set (100 participants, 121 parameter combinations) we also examined whether the resulting behaviour could be better fit using a random model. In general, model recovery was very good – for most parameter combinations, most of the simulated data sets were better fit with an RL model compared to a random model. We noted that a random model occasionally provided a better fit when the learning rate was low ($\alpha$ = 0.01) – See bottom of Supplementary Figure 3. We therefore restricted our model fitting procedure to only test $\alpha$'s from 0.1 to 1.

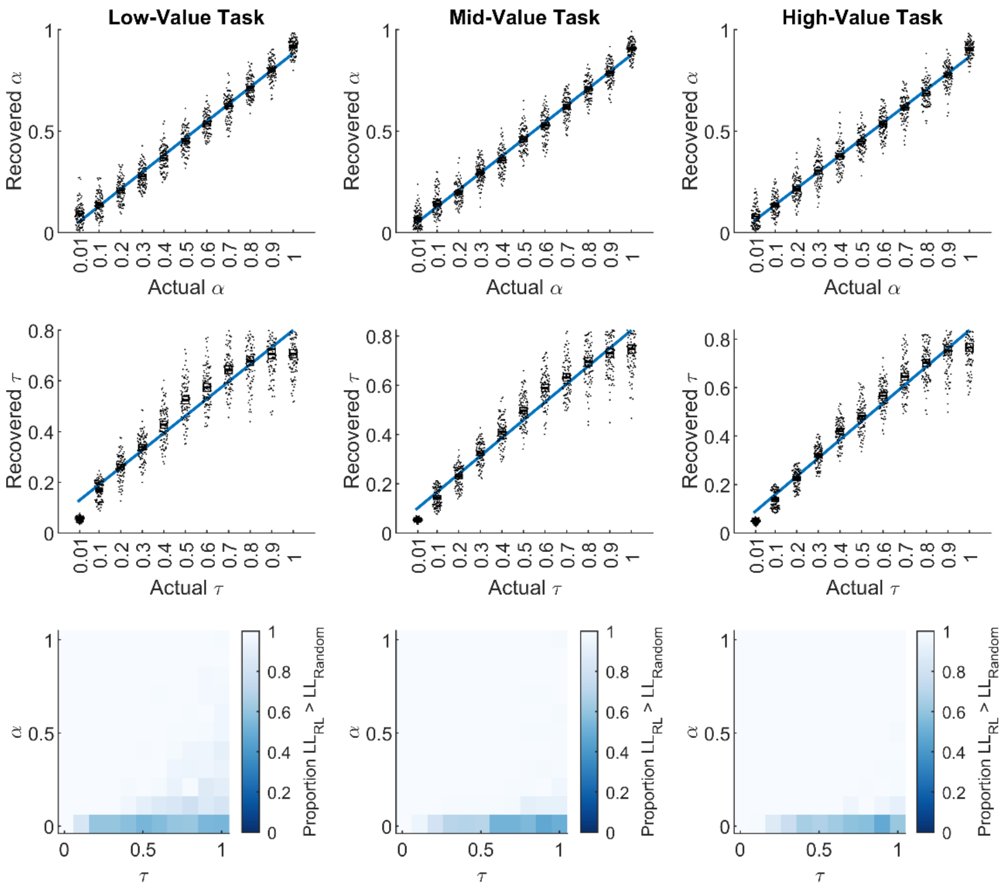


*Supplementary Figure 3.* **Parameters and model were recoverable.** Both learning rate (top row) and temperature (middle row) were recoverable following simulation with a reinforcement learning model. For most parameter combinations, a reinforcement learning model provided a much better fit to simulated data compared to a random model (100 simulated participants per combination, bottom row).

### Time-Frequency Analysis

While investigating the cue-locked response, we observed what appeared to be differences in oscillatory activity. We then conducted an exploratory analysis examining frontal midline theta (FMT), which consists of oscillatory power in the 3–8 Hz range over frontal-central scalp locations. FMT (or FMΘ) is thought to originate in ACC and FMT is thought to index a cognitive control-related process (Cavanagh et al., 2012; Cavanagh & Frank, 2014). A variety of events have been shown to elicit increased FMT power including loss feedback (Li et al., 2018) and predictive cues (Cavanagh et al., 2011). Like the RewP, FMT provides a readout of ACC activity, but one that is dissociable from the RewP (Hajihosseini & Holroyd, 2013; Holroyd & Umemoto, 2016; Paul et al., 2020). In particular, whereas the RewP reflects mainly phase-locked activity, FMT can reveal both phase-locked and non-phased-locked activity (Cohen et al., 2008).

To quantify FMT in response to cues, we first created cue-locked epochs from 600 ms pre-cue to 1200 ms post-cue. Compared to our ERP analysis, longer epochs were necessary to account for potential edge artifacts associated with time-frequency analysis. Using the same artifact rejection criteria as before, we excluded on average 2.46%, 95% CI [1.06, 3.86] from analysis. We then transformed each cue-locked EEG epoch into its time-frequency representation. This was done by convolving the EEG with wavelets of the form $e^{i2\pi ft}e^{-t^2/2\sigma^2}$ for frequencies *f* ranging from 1 to 30 in 60 logarithmic steps. The parameter $\sigma$, which affects wavelet width, was set to $6/2\pi f$. The convolution was squared to compute power *p(t)*, averaged for each participant and condition, and converted to the decibel scale according to the equation $10\log_{10}(\frac{p(t)}{p(baseline)})$. The baseline, *p(baseline)*, was defined as the mean power from -400 to -100 ms relative to the event of interest (cue or loss feedback). Grand averages were constructed for each condition by averaging time-frequency data across participants.

We focused on electrode FCz, in line with previous work linking FMT with feedback processing and cognitive control (Cavanagh et al., 2012; Williams et al., 2021). As this was an exploratory analysis, we first collapsed the data over all cue conditions and then selected power values greater than 0, for all times within 3-8 Hz, corresponding to the frequency range used in previous studies (Cavanagh et al., 2012; Williams et al., 2021). We then computed the mean power within this cluster for each condition and participant. See Supplementary Figure 4 for the collapsed mean time-frequency response.

To determine whether cues of different value elicited different adjustments to cognitive control, we compared FMT using repeated-measures t-tests. This was done for the task in which both cue types appeared (the mid-value task). We also compared cue-locked FMT to low-value cues in the low-value task to high-value cues in the high-value task. No other comparisons were made as this would require comparing events of differing frequency of occurrence, e.g., low-value cues were more frequent in the low-value task compared to the mid-value task. This is important because, like the RewP, FMT is sensitive to probability-related expectations (Cavanagh et al., 2012).

#### *Results*

There was no cue-locked FMT difference between low-value (0.22 dB, 95% CI [-0.14, 0.57]]) and high-value (0.15, 95% CI [-0.24, 0.54]) cues in the mid-value task, $t(24) = 0.45$, $p = .66$, Cohen's $d = 0.09$. There was also no cue-locked FMT difference between low-value cues in the low-value task (0.29 dB, 95% CI [-0.05, 0.63]) and high-value cues in the high-value task (0.31, 95% CI [-0.10, 0.73]), $t(24) = -0.21$, $p = .84$. Cohen's $d = -0.04$. See Supplementary Figure 5.

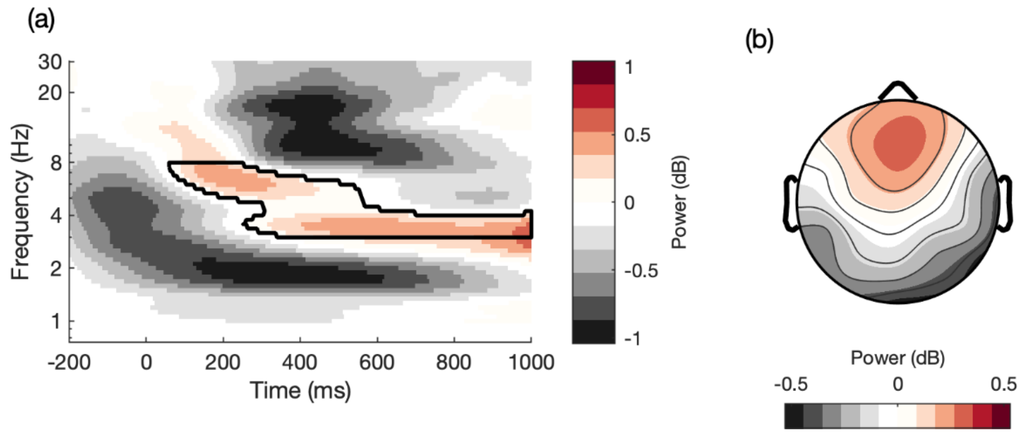


*Supplementary Figure 4.* **Mean cue-locked FMT.** (a) Time-frequency response collapsed across all cue conditions. The black outline shows the region of analysis. (b) Scalp distribution of the mean time-frequency response.

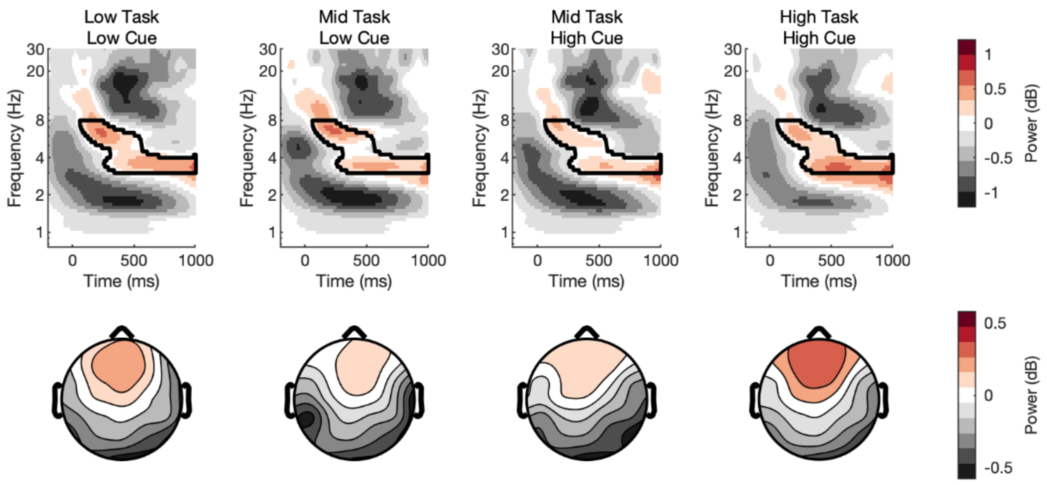


*Supplementary Figure 5.* **No cue-locked FMT differences.** Time-frequency plots at FCz and scalp topographies for EEG locked to cues in each task (task value: low, mid, high; cue value: low, high). Cue-locked FMT did not differ by cue value.

We then examined whether there was a relationship between FMT and performance. Because the goal of this analysis was to assess individual differences, the data of all of the participants (learners and non-learners) were included. In the mid-value task, we observed that performance correlated with FMT to low-value cues, $r(33) = 0.37$, $p = .03$ but not high-value cues, $r(33) = 0.22$, $p = .19$ (Supplementary Figure 4a–b). In the high-value task, we observed that performance correlated with FMT to high-value cues, $r(33) = 0.49$, $p = 0.003$ (Supplementary Figure 4c).

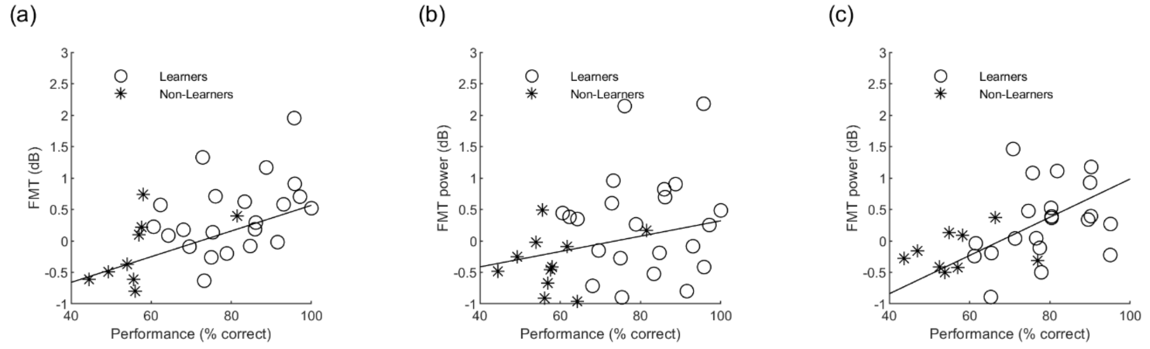


*Supplementary Figure 6.* **Cue-locked frontal midline theta (FMT) depends on cue and task value.** Performance in the mid-value task correlated positively with low-value cue FMT (a) but not high-value cue FMT (b). (c) The relationship between cue-locked FMT and performance was also present in the high-value task.
